# Characteristics of Beta Waveform Shape in Parkinson’s Disease Detected with Scalp Electroencephalography

**DOI:** 10.1101/534396

**Authors:** Nicko Jackson, Scott R. Cole, Bradley Voytek, Nicole C. Swann

**Author notes:** Please address correspondence to: Dr. Nicole C. Swann, 181 Esslinger Hall, 1525 University St, Eugene, OR 97403.

## Abstract

Neural activity in the beta frequency range (13-30 Hz) is excessively synchronized in Parkinson’s Disease (PD). Previous work using invasive intracranial recordings and non-invasive scalp electroencephalography (EEG) has shown that correlations between beta phase and broad-band gamma amplitude (i.e., phase amplitude coupling) are elevated in PD, perhaps a reflection of this synchrony. Recently, it has also been shown, in invasive human recordings, that nonsinusoidal features of beta oscillation shape also characterize PD. Here we show that these features of beta waveform shape also distinguish PD patients on and off medication using non-invasive recordings in a dataset of 15 PD patients with resting scalp EEG. Specifically, beta oscillations over sensorimotor electrodes in PD patients off medication had greater sharpness asymmetry and steepness asymmetry than on medication (sign rank, p=0.006, p=0.003 respectively). We also showed that beta oscillations over sensorimotor cortex most often had a canonical shape, and that using this prototypical shape as an inclusion-criteria increased the effect size of our findings. Together our findings suggest that novel ways of measuring beta synchrony that incorporate waveform shape could improve detection of PD pathophysiology in non-invasive recordings.

## Introduction

Parkinson’s disease (PD) is characterized by excessively synchronized neural activity in the beta frequency range (13-30 Hz)(Brown and Williams, 2005; Brown, 2007; Moran et al., 2008). Much of the work describing this phenomenon has come from invasive single unit and local field potential recordings from the human basal ganglia. Together this body of work has formed a compelling narrative wherein motor system beta synchrony, which can be detected using traditional beta power in basal ganglia, is elevated in PD patients off medication and reduced by therapies like medication and deep brain stimulation (DBS)(Kühn et al., 2006; Brown, 2007; Kühn et al., 2008). This has motivated the idea that measures of brain physiology that reflect beta synchrony could be potential “biomarkers” for PD pathophysiology. Such objective measures of PD symptoms would have enormous clinical potential for diagnosis, monitoring, and adjusting patient therapies. However, basal ganglia recordings can be difficult to acquire in humans and necessitate invasive approaches. Therefore, a cortical signature, which could be acquired non-invasively, or less invasively, could provide greater clinical utility.

While conventional signal processing measures, such as beta power, have failed to reliably differentiate PD as a function of severity or diagnosis at the cortex (Stoffers et al., 2007; de Hemptinne et al., 2013; George et al., 2013; de Hemptinne et al., 2015; Swann et al., 2015; Malekmohammadi et al., 2018), novel metrics such as phase amplitude coupling between beta and broadband gamma (50-150 Hz) (PAC) have proven more promising. Specifically, PAC over motor cortex detected using electrocorticography (ECoG) is elevated in PD compared to other groups (de Hemptinne et al., 2013) and is reduced with DBS in a clinically-relevant manner (de Hemptinne et al., 2015; Malekmohammadi et al., 2018). Interestingly, following characterization of PAC with ECoG, it was demonstrated that elevated PAC can also be detected non-invasively with scalp electroencephalography (EEG) (Swann et al., 2015). Moreover, PAC recorded with scalp EEG could differentiate PD patients on and off medication and differentiate PD patients off medication from healthy controls, at the group level.

Recently, increased attention has been given to novel waveform metrics, that may provide additional insights into neurophysiological mechanisms (Cole and Voytek, 2017). Using ECoG recordings, recent work has shown that beta waveform shape, similar to PAC, differentiates PD patients on and off DBS (Cole et al., 2017). Specifically, PD patients off DBS had more sharp and asymmetric beta activity, which was reduced on DBS. These waveform metrics may provide novel insights into underlying PD pathophysiology. Particularly, cortical beta waveform shape may indicate the aggregate of synchronous inputs (perhaps from the basal ganglia via the thalamus) onto cortical pyramidal neurons (Sherman et al., 2016; Cole and Voytek, 2018a).

Given the potential utility of considering waveform shape, and the past literature that demonstrated that signatures reflecting PD pathophysiology initially characterized with ECoG are also detectable with EEG, we sought to investigate if waveform shape detected with scalp EEG might also be an electrophysiological biomarker for PD pathophysiology. Specifically, we tested if the shape of sensorimotor beta activity in scalp EEG differed between PD patients on and off medication and PD patients compared to healthy age-matched control participants. We were especially interested in investigating how beta waveform shape compares to PAC in its ability to distinguish medication state and what the strengths and weaknesses might be for each measure as a neurophysiological biomarker.

## Methods

### Participants

We analyzed a previously published data set, from another laboratory (George et al., 2013). This same data was analyzed in another published report which showed elevated PAC between beta phase and broadband gamma amplitude in PD patients off medication compared to the same patients on medication and compared to a group of healthy, age-matched, control participants (Swann et al., 2015). This dataset includes EEG data from 15 PD patients (8 female, mean age = 63.2+/-8.2 years) on and off dopaminergic medication and 16 healthy, age-matched, control participants (9 female, mean age, 63.5 +/-9.6 years). All PD patients had been diagnosed by a movement disorder specialist at Scripps clinic in La Jolla, California. Participants were right-handed and provided written consent in accordance to the IRB of the University of California, San Diego and the Declaration of Helsinki. For additional patient information, see (George et al., 2013).

### Data collection

Data from patients on and off medication were collected on different days with a counterbalanced order. For the on medication recordings, patients continued their typical medication regimen. For the off medication state, patients discontinued medication use at least 12 hours before the session. Control participants were tested once. EEG data were acquired using a 32-channel BioSemi ActiveTwo system, sampled at 512 Hz. Resting data were recorded for at least 3 minutes. During data collection, the participants were seated comfortably and told to fixate on a cross presented on a screen. Additional electrodes were placed lateral to and below the left eye to monitor eye blinks and movements. Participants also completed several other assessments described in the previously published report (George et al., 2013), which were not analyzed here.

### Data preprocessing

Data were preprocessed in MATLAB using custom scripts and EEGLAB functions (Delorme and Makeig, 2004). In brief, first, each electrode’s mean was removed and then a common average reference was applied. Excessively noisy electrodes were excluded. The data were then high pass filtered at 0.5 Hz to remove low frequency drift (using a 2 way FIR filter, eegfilt (Delorme and Makeig, 2004). Then, data were manually examined for artifacts (eye blinks and movements, muscle activity, electrical noise, and other sources of noise), and indices containing these events were flagged for rejection. In order to avoid filtering over non-continuous data, rejection regions were excluded after filtering (as described below). In addition to common average reference, we also used a current source density (CSD) approach (CSDtoolbox in MATLAB, spline flexibility = 3, (Kayser, 2009)) for an auxiliary analysis to compare the current results to a previously published report (Swann et al., 2015).

We focused predominantly on electrodes closest to sensorimotor cortex (C3 and C4). To compare data from these electrodes to recordings more likely to be impacted by residual electromyogram (EMG) artifact we also examined the electrodes closest to the temporalis muscles (F7 and F8) (Swann et al., 2015). Unless otherwise specified, statistical testing focused on a “composite” sensorimotor signal where measures from C3 and C4 were calculated separately and then results were averaged for each participant, such that each participant contributed one data point, prior to performing statistical testing across participants. The same procedure was used for electrodes closest to the temporalis muscles.

### Data Analysis

The following waveform metrics were calculated using Python code that were modified from (Cole et al., 2017), unless otherwise specified. The original code can be found at https://github.com/voytekresearch/Cole_2017. The code we used for this manuscript can be found at https://github.com/SwannLab/Jackson_2019.

### Sharpness Ratio

We calculated sharpness ratio using previously defined methods (Cole et al., 2017). In brief, each signal was filtered between 13-30 Hz using a window-method based FIR filter (scipy.filter.firwin, order: 231 ms). Indices of rising and falling zerocrossings (i.e. voltage sign-changes) were identified in this filtered signal. Next, returning to the raw signal, the indices of the maximum voltage (the ‘peak’), and minimum voltage (the ‘trough’), between zerocrossings were found. Once peaks and troughs were identified, those that fell in regions previously marked for rejection due to artifacts were excluded from further analysis. Peak sharpness was defined as the mean of the voltage difference between the peak and 3 data points before and after the peak. Trough sharpness was calculated in an analogous way. Three sample points corresponds to ∼5.9 ms of data. Similar to a previous report (Cole et al., 2017), calculating sharpness ratio using different widths produced similar results. Sharpness ratio was defined as the log of the maximum ratio of either mean peak sharpness to mean trough sharpness, or mean trough sharpness to mean peak sharpness (Cole et al., 2017). The maximum was taken so that logged ratios were positive. See Figure 1 for schematic explaining experimental methods.

**Figure 1.**
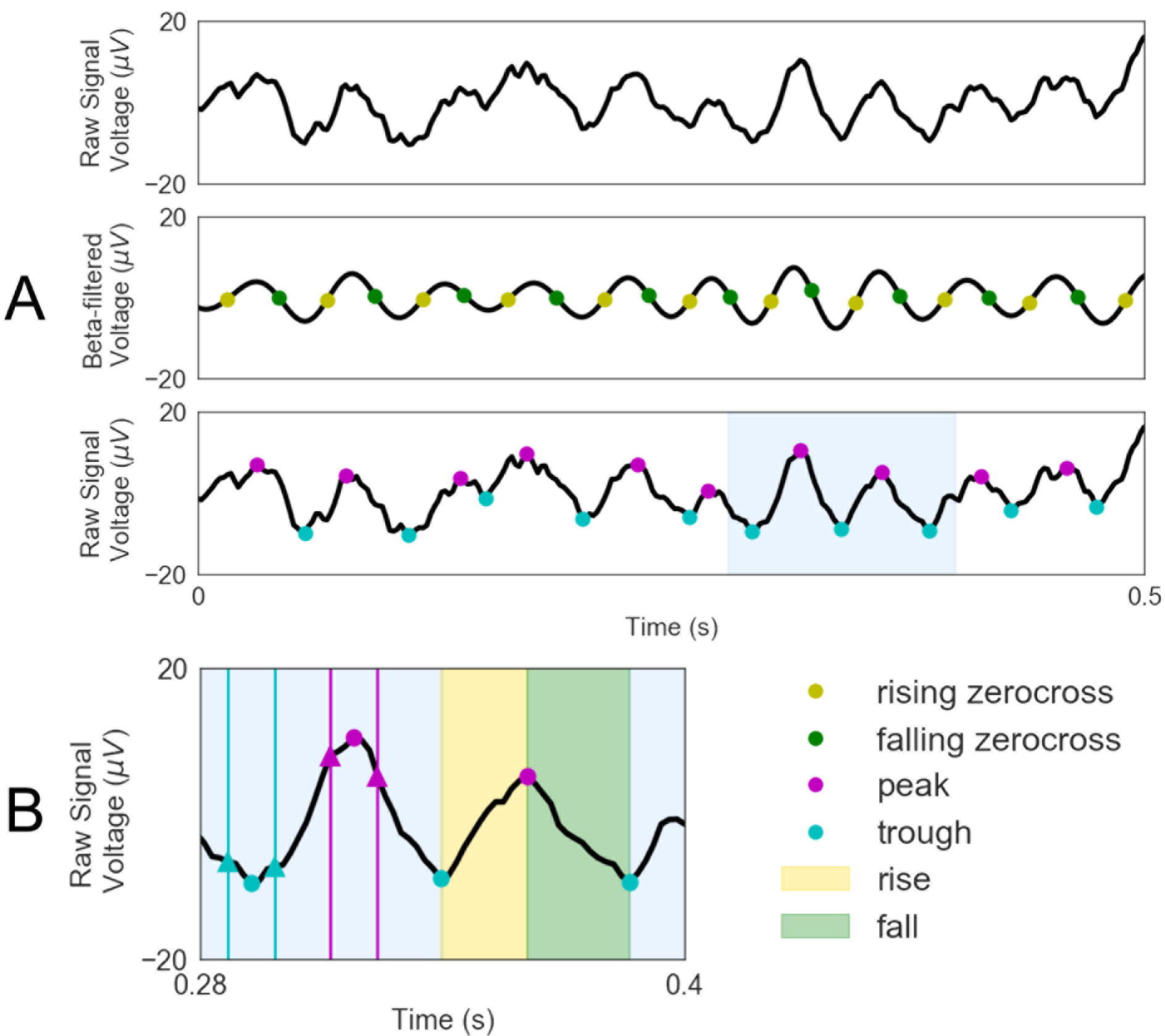
Schematic summarizing waveform shape calculations. A) First rising and falling zerocrossings were identified from the beta filtered signal. Then, peaks and troughs that lie between adjacent zerocrossings were found in the raw signal. B) Peak sharpness was calculated as the mean of the difference between the peak (pink circle) and the voltage points 3 samples before and after the peak (pink triangles). The trough sharpness was calculated in a similar fashion. The yellow area, between a trough and subsequent peak, indicates an area where rise steepness is determined, while the green area, between a peak and subsequent trough, indicates an area where decay steepness was found. Figure modeled after (Cole et al., 2017), but here is shown with the present EEG data.

**Figure 2.**
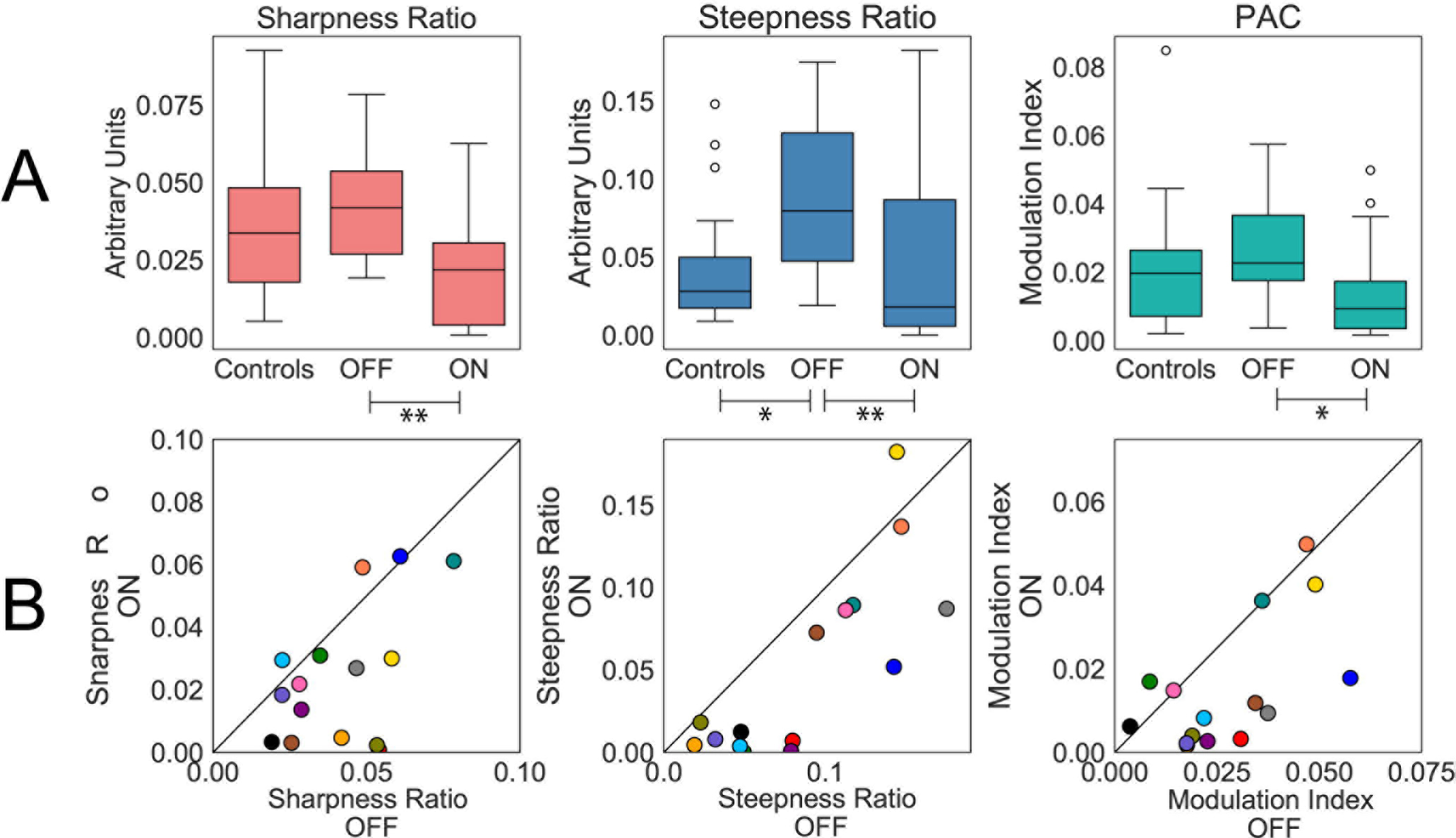
Sharpness Ratio, Steepness Ratio, and PAC all decreased with medication in PD patients. A) Box plots of each measure for patients off medication, on medication, and control participants. Double asterisks represent significance at p < 0.01 and a single asterisk represents significance at p < 0.05 B) Individual data for each measure and each patient on and off medications. Each color corresponds to a different participant. Diagonal lines represent unity.

### Steepness Ratio

Steepness ratio was also calculated according to previously published methods (Cole et al., 2017). Rise steepness, between a trough and a subsequent peak, was defined as the maximum value of the first derivative, or the greatest slope, of the signal. Similarly, decay steepness was defined as the maximum of the absolute value of the slope between each peak and subsequent trough. Steepness ratio was calculated as either the ratio of mean decay steepness to mean rise steepness or vice versa, such that the logged ratio was positive.

### Phase Amplitude Coupling

Phase Amplitude Coupling (PAC), which reflects correlations between beta phase and broad-band gamma amplitude, was calculated using the normalized modulation index metric (Ozkurt and Schnitzler, 2011), with the open-source package, pacpy, which can be found at https://github.com/voytekresearch/pacpy. In brief, using a FIR filter, beta and gamma components were extracted from the signal. Beta was filtered between 13 and 30 Hz with a filter order of 231ms. Gamma was filtered between 50 and 150 Hz with a filter order of 240 ms. After filtering, regions that had been marked as containing artifacts were removed. Beta phase and gamma amplitude time series were calculated by extracting the angle and amplitude of the Hilbert transform respectively. Note that PAC results have already been published using these data (Swann et al., 2015). However, we repeat this analysis here for direct comparison with the waveform shape metrics. Note also that here we used an alternative method to calculate PAC to be consistent with previous comparisons between waveform shape and PAC (Cole et al., 2017). However, results were similar between methods, except where noted. Here, we also used a different referencing scheme (average reference compared to CSD in the previous report).

### Peak-to-trough Ratio and Rise-to-fall Ratio

Sharpness ratio and steepness ratio quantify overall asymmetry but do not characterize the specific peak/trough or rise/fall contributions to this asymmetry. To address this, and further characterize waveform shape, we calculated peak-to-trough ratio (analogous to sharpness) and rise-to-fall ratio (analogous to steepness). Peak-to-trough ratio is the log of mean peak sharpness divided by mean trough sharpness, so that a positive peak-to-trough ratio indicates relatively sharper peaks while a negative ratio indicates relatively sharper troughs. Similarly, rise-to-fall ratio is the log of the mean rise steepness divided by the mean fall steepness, and a positive rise-to-fall ratio signifies a relatively steeper rise and a negative rise-to-fall ratio indicates a relatively steeper decay. Because these measures differentiate peaks/troughs and rise/fall, they allow typification of waveform shape into four distinct categories (i.e. negative vs positive peak-to-trough ratio and rise-to-fall ratios, Figure 4C).

**Figure 3.**
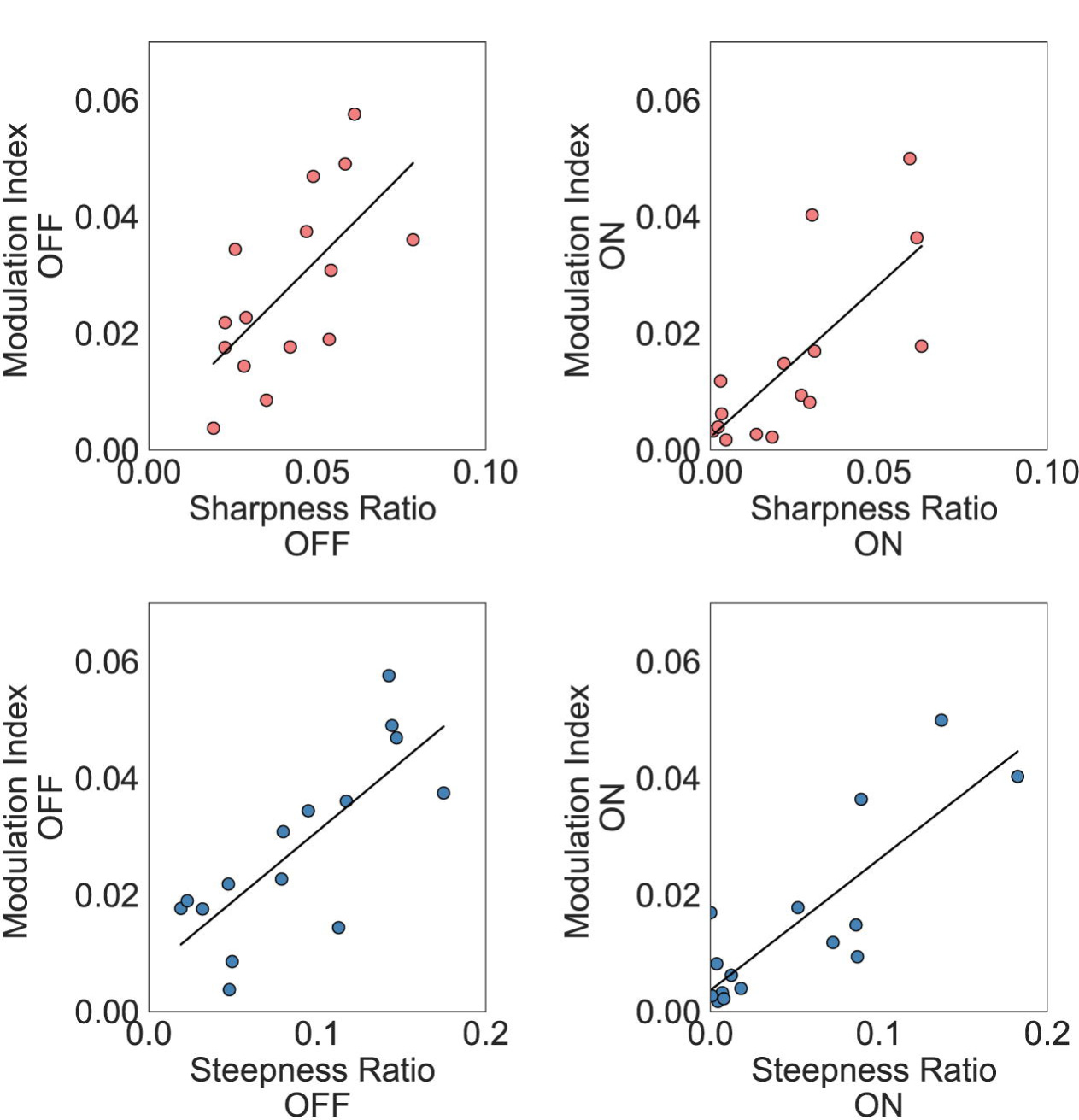
Both sharpness ratio and steepness ratio over sensorimotor cortex correlate with PAC in both medication states.

**Figure 4.**
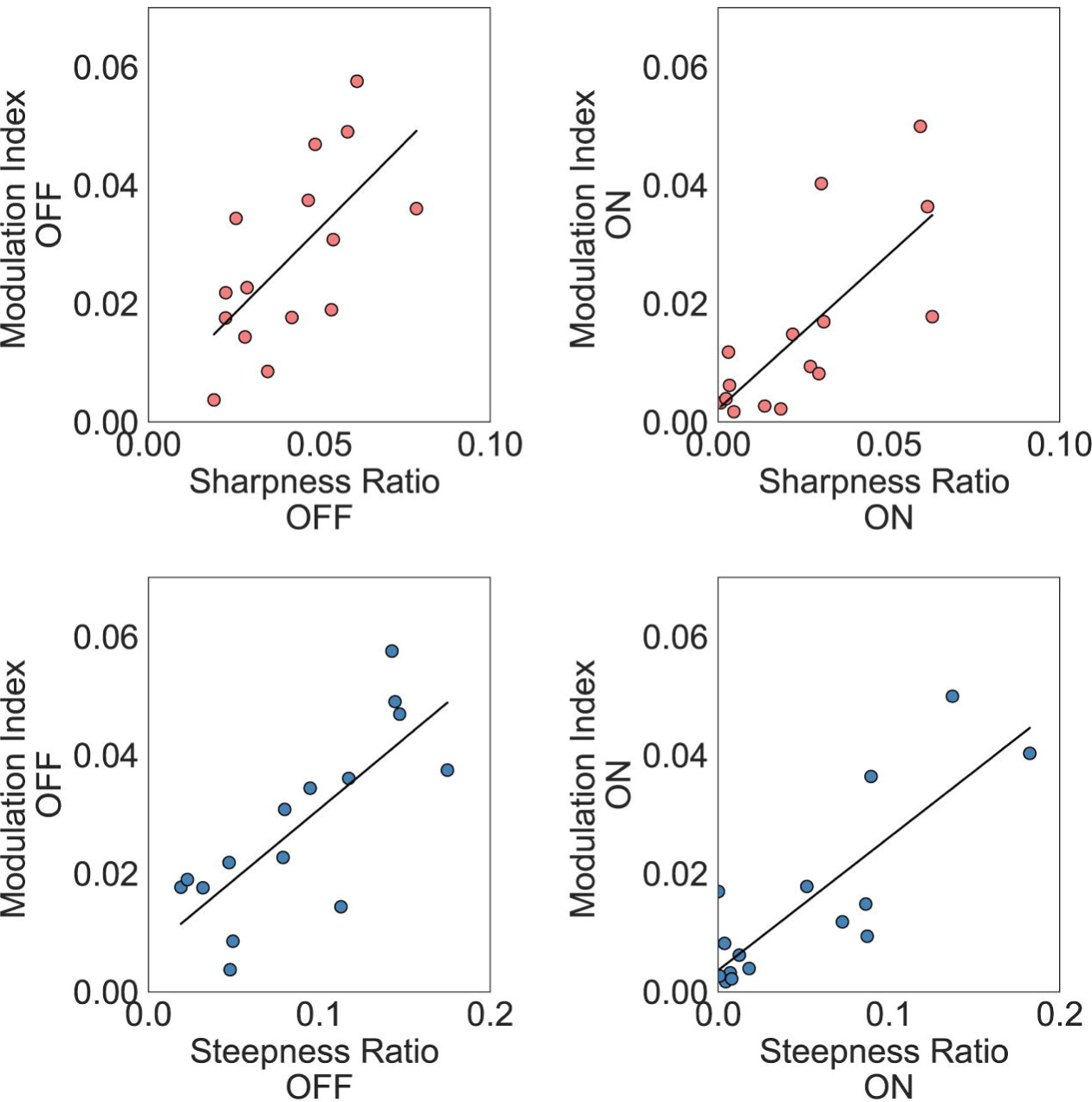
Sensorimotor cortex recordings have a canonical shape. A. Peak-to-trough ratio versus rise-to-fall ratio over sensorimotor electrodes (C3 and C4) shown with individual participants/electrodes. Circles represent off medication, while squares represent on medication. The color map corresponds to PAC values. Only patient data is shown, but control data followed a similar pattern. B. Peak-to-trough versus rise-to-fall ratio in electrodes closest to temporalis muscles (F7 and F8) shown separately. Note the scale of the PAC color map is different than in A. C. Representative waveform shape for each quadrant. The sensorimotor data falls mainly in quadrant 4 (blue). This corresponds to sharper peaks and steeper decays.

### Analysis over time

For most measures, sharpness ratio, steepness ratio, and PAC were calculated for the entire resting-state recording. The exception was the analysis shown in Figure 5 that demonstrates variability over time for each measure. Here sharpness ratio, steepness ratio, and PAC were calculated as previously described above except using 10 second non-overlapping windows for each patient individually on and off medication. Using this approach, we assessed intra-subject variability by looking at values for all windows over time at the individual participant level.

**Figure 5.**
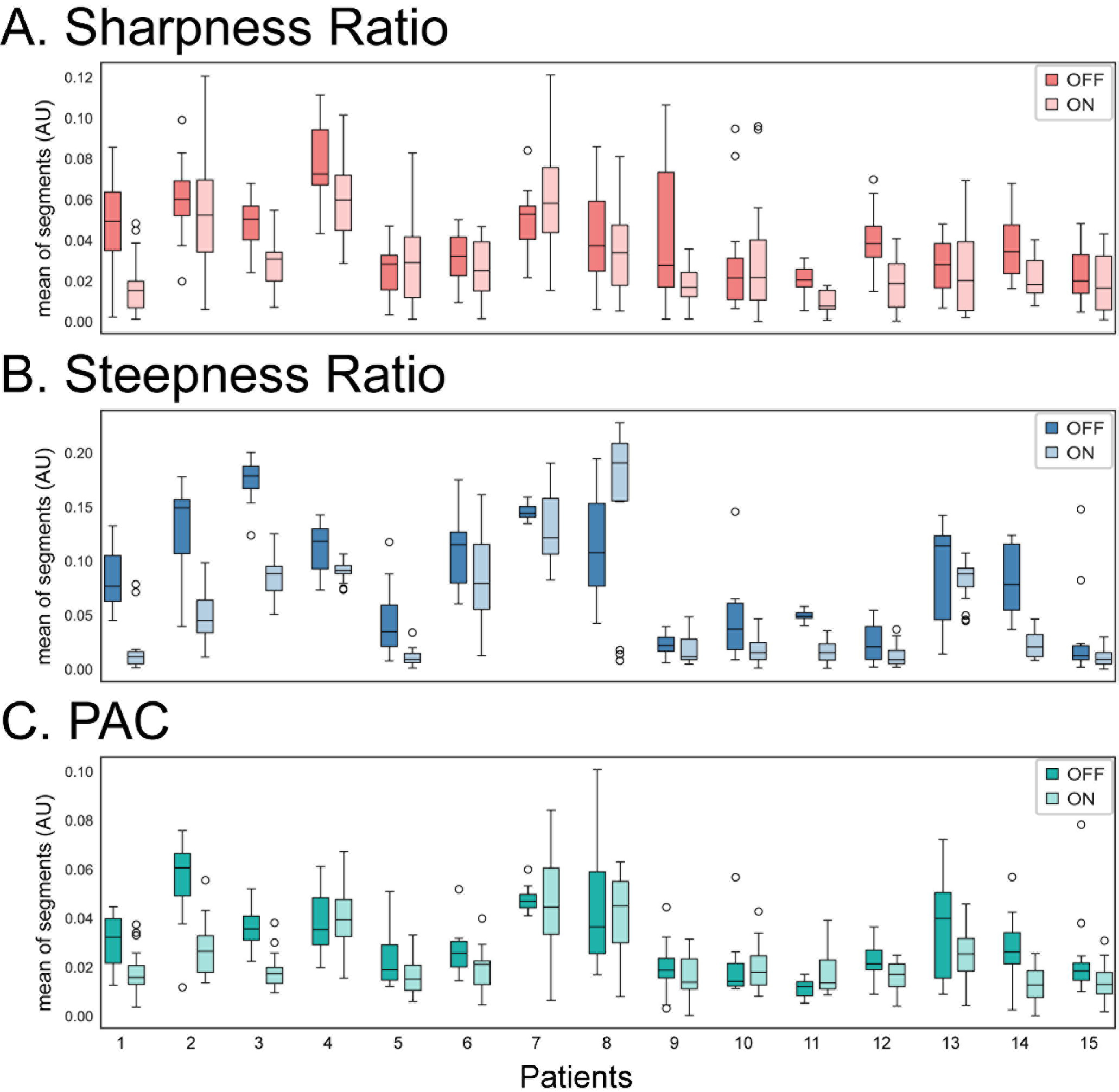
Waveform metrics and PAC change over time. Each plot shows the distribution of each measure calculated over 10 second non-overlapping segments for each patient on and off medication.

### Statistical tests

For across group comparisons we used a non-parametric Wilcoxon signed rank test for all comparisons of patients on and off medication and Wilcoxon rank sum test to compare patients off medication to controls. Spearman correlation coefficients were calculated to compare similarities between PAC and waveform shape-based metrics.

We also calculated effect size for each waveform metric. Effect size (Cohen’s d) quantitatively measures the magnitude of an effect (Cohen, 1998). For these calculations, PAC was log scaled in order to ensure normalization. Sharpness ratio and steepness ratio were already log scaled as part of their standard analysis.

## Results

### Waveform metrics differentiate PD medication state

Medication reduced sharpness ratio (Figure 2; p=0.006) and steepness ratio (Figure 2; p=0.003) in PD patients over sensorimotor electrodes. Notably, sharpness ratio and steepness ratio are higher for waves with asymmetrical waveforms (e.g. sawtooth or arched waves), suggesting that medication flattened waveforms or made them more symmetric. PAC also decreased with medication (Figure 2; p=0.011), as has been previously published with this same data (but using an alternative method to calculate PAC and an alternative referencing scheme.)

Data from electrodes closest to the temporalis muscles (F7 and F8) were also examined to compare the sensorimotor results to electrodes more likely to be contaminated by muscle activity and address our concerns that differences in muscle activity may have been contributing to differences. However, none of these measures differed between patients on and off medication over these electrodes: sharpness ratio, p=0.73, steepness ratio, p=0.394, PAC, p=0.191 (as was previously published, using an alternative PAC calculation and alternative referencing scheme).

We also examined PD patients off medication and control participants. There was no significance difference between these groups for sharpness ratio (p=0.494), however steepness ratio did distinguish patients from controls (p=0.020). Somewhat surprisingly there was also no significant difference between PD patients off medication and control participants for the PAC calculation (p=0.236). However, a previous publication testing these same data, but using a CSD referencing scheme (which is designed to increase spatial specificity), and an alternative method for calculating PAC (Tort et al., 2008) did find a significant difference between PD patients off medication and control participants (Swann et al., 2015). When we tested the CSD density referencing scheme using the current method for PAC calculation, there was a significant difference between patients off medication and controls (p=0.022), confirming findings from the previous publication which suggest PAC differences between patients and controls were sensitive to referencing scheme and were most apparent for approaches which isolate spatial sources (Swann et al., 2015). Note that we chose not to use the CSD reference for the waveform shape calculations since this scheme might cause unpredictable phase inversions.

Overall, the difference between patients on and off medication was robust over sensorimotor cortex with significant differences for each of our measures. In contrast, measures over electrodes more likely to be contaminated by electromyogram artifact (i.e. closest to temporalis muscles) did not differentiate groups for any measure. Therefore, we believe that our findings are less likely to be driven by muscle artifact.

The comparison between patients off medication and controls was less clear with only steepness ratio distinguishing the groups (and PAC calculated using a CSD reference as was previously reported (Swann et al., 2015)).

### Relationships between Sharpness/Steepness Ratio and PAC

To address how waveform shape and PAC might relate to one another and probe if they may be reflecting the same underlying process or picking up on different aspects of pathophysiology we examined relationships between the waveform shape measures and PAC. Like the previous report using invasive recordings (Cole et al., 2017) sharpness ratio and PAC were correlated in both on and off medication states in sensorimotor electrodes (Figure 3. Off medication: r = 0.70, p=0.004; on medication: r = 0.74, p=0.002). Likewise, steepness ratio and PAC are also highly correlated for both medication states (off medication: r = 0.74, p=0.002; on medication: r = 0.68, p=0.006).

### Waveform shape

We observed a negative correlation between the peak-to-trough and rise-to-fall ratios of the oscillations (spearman, r = −0.73, p=4e-11). In general, sensorimotor electrode oscillations had a sharper peak with a steeper decay, falling in quadrant 4 (Q4). Arch shapes (sometimes referred to as a “mu” shape) have previously been described in sensorimotor recordings in this frequency range (Figure 4A) (Pfurtscheller et al., 1997). In contrast, electrodes closest to temporalis muscles had a more variable waveform shape pattern, perhaps consistent with contributions from both EEG and EMG, and/or the general lack of a consistent dominant waveform pattern (Figure 4B). Consistent with the above figures, waveform shape is more symmetrical when PD patients were on medication, as indicated by their central location on the following figures. Furthermore, Figure 4A shows that as sharpness ratio and steepness ratio approach zero, PAC decreases (i.e. lower PAC values are closer to the origin and vice versa) as expected from the correlations described above (Figure 3).

We next tested if including only sensorimotor data that fell in Q4 would strengthen medication state differentiation (since this quadrant captured the dominant shape of the sensorimotor activity). The rationale is that if this shape is a canonical feature of sensorimotor EEG data, using it as an inclusion-criteria might exclude periods of time when beta power is lower, leading to unreliable measures, and periods where residual EMG may be obscuring activity. For this analysis, patients were only included if they had data on and off medication which fell in Q4. Out of 15 patients, 9 patients met this criteria for C3 and 8 met the criteria for C4. When data from both C3 and C4 fell within Q4 (8 patients), measures were calculated for each electrode separately and then averaged so that each patient contributed 1 data point for the effect size calculation, analogous to the above analyses. When only 1 electrode was available (1 patient), measures from just this electrode were used for the effect size analysis. Using this approach there were 9 total Q4 observations for each waveform metric. We compared effect size when all data were used for the calculation, to the effect size when only data from Q4 were included. Effect size increased for all measures, at least marginally, when using the Q4 shape as an inclusion criteria (Table 1)

**Table 1.** Effect size (Cohen’s d) for all patients and for quadrant 4 (Q4) patients, on versus off medication. All metrics were log scaled.

| Metric | All | Q4 |
| --- | --- | --- |
| Sharpness ratio | 0.86 | 0.95 |
| Steepness ratio | 0.68 | 0.70 |
| PAC | 0.81 | 1.07 |

### Intra-subject Variability

To investigate how much these signals fluctuate over time, even at rest, we examined values of each metric for small segments of data (10 seconds) over a longer recording (Figure 5). This approach shows that these signals are not static, and in fact vary quite a bit for some participants, suggesting that the amount of data needed to see differences may be participant dependent. To quantify this variability, we calculated the effect size of the on medication versus off medication comparison for each patient individually. Using this approach the average and standard deviation (respectively) of effect size was 0.68 and 0.73 for sharpness ratio, 1.53 and 1.50 steepness ratio, and 0.66 and 0.81 for PAC. Thus, there was significant variability in the robustness of these effects between participants.

## Discussion

Converging work has shown that PD is associated with excessive beta synchrony. Here we show that this pathophysiological synchrony is manifest in a change in waveform shape which can be detected at the scalp. Specifically, we show beta waveforms are more asymmetric for PD patients off medication and that this is reduced with medication. Further, we show that including only data with a canonical arched waveform shape, consistent with a sensorimotor “mu” shape, increased effect size for all metrics examined. This suggests the consideration of waveform shape could improve the efficacy of non-invasive PD biomarkers. Importantly, this difference in waveform shape occurred in the context of a lack of a consistent difference in conventional spectral power in the beta range over sensorimotor cortex in previous publications examining PD (Stoffers et al., 2007; de Hemptinne et al., 2013; de Hemptinne et al., 2015; Malekmohammadi et al., 2018), including two which analyzed the same dataset we examined here (George et al., 2013; Swann et al., 2015). This underscores the general importance of including novel metrics, such as waveform shape, in analyses of electrophysiological data, including scalp EEG.

### Mechanisms of waveform shape asymmetry and PAC

Converging evidence suggests that PAC is elevated in untreated PD patients compared to treated (de Hemptinne et al., 2015; Swann et al., 2015; Malekmohammadi et al., 2018). It also emerges during the induction of severe parkinsonian symptoms in non-human primate models, and is related to disease severity (Devergnas et al., 2017). However, the etiology of PAC is unclear. One potential mechanism of PAC is that it describes a relationship between lower frequency activity and high frequency broadband amplitude, a putative surrogate for spiking (Manning et al., 2009). Thus, synchronous cortical input in beta (perhaps from basal ganglia via thalamus) may bias the probability of neural spiking. This is supported in PD, by the observations of unit spiking coupled to beta local field potentials in basal ganglia recordings (Moran et al., 2008). However, several other phenomena can also lead to increased PAC, including periodic sharpness or transients (Kramer et al., 2008; Aru et al., 2015). Comparably, sensorimotor waveform shape may reflect synchronous inputs (Sherman et al., 2016) perhaps from basal ganglia (Nini et al., 1995; Sharott et al., 2005; Brown, 2007; Mallet et al., 2008), that are not necessarily associated with spiking. Excessive basal ganglia-thalamocortical loop synchrony may constrain neurons in an inflexible pattern, prevent changes necessary for dynamic behavior, and effectively hinder neural communication by exhausting ‘neural bandwidth’ (Reyes, 2003; de Hemptinne et al., 2013; Swann et al., 2015; Voytek and Knight, 2015; Cole et al., 2017). We suspect that waveform shape metrics and PAC (and likely other measures elevated in untreated PD such as global beta coherence across electrodes (Silberstein et al., 2005)) are imperfect ways of measuring the same underlying pathophysiology - excessive beta synchrony and neural entrainment. This synchrony exists in the healthy state, but is amplified throughout motor networks, both within and between individual structures in PD. Dopaminergic medication reduces this excessive synchrony, in conjunction with improvement of the Parkinsonian state, which is reflected in the reduction of PAC, sharpness ratio, and steepness ratio (de Hemptinne et al., 2015).

### Differences and similarities between measures

Consistent with previous findings in ECOG recordings (Cole et al., 2017), waveform shape (sharpness ratio and steepness ratio) is correlated with PAC, indicating that they may be picking up similar characteristics. Indeed, Cole and colleagues showed remarkable correspondence between broadband amplitude measured in PAC and nonsinusoidal aspects of waveform shape. Therefore, it could be that elevated PAC may be better described as a change in waveform shape, a more parsimonious explanation of the signal (i.e. a measure of one periodic process – beta, rather than two – beta and gamma)(Cole et al., 2017). Alternatively (or additionally) PAC and waveform shape may be correlated because they both relate to PD severity or PD pathophysiology.

Despite high correlations between measures, PAC could not be completely explained by waveform shape in our data, and the measures show differences in detection ability, especially for PD patients off medications compared to healthy controls. This suggests that different measures may have different strengths and weaknesses, even if detecting the same underlying pathophysiology.

### Possible ways to improve the electrophysiology biomarker

Considering waveform shape as an inclusion-criteria for our group analyses improved the effect size, suggesting that development of a method to extract waveforms of a specific shape from a signal might result in a more powerful biomarker. This may be especially important for EEG data, which is more likely to suffer from lower signal-to-noise ratio. Therefore, developing a method to select only data with specific shapes, perhaps by building a filter which only extracts signals with a canonical sensorimotor shape (i.e. a sharp peak and steep fall), might improve classification since it would isolate data with prototypical patterns consistent with a sensorimotor origin. Additionally, this filter could also be used to detect oscillatory bursts with this shape within the beta frequency range (Tinkhauser et al., 2017) or to index the cycle-to-cycle variation in a beta signal (Cole and Voytek, 2018b; Torrecillos et al., 2018). Alternatively, principal component analysis could be used to create an optimal waveform shape metric that is comprised of weighted components of each waveform meausure.

### Changes of the measures with behavior and spontaneous fluctuations

Previous work has shown that PAC fluctuates during movement (Miller et al., 2012; de Hemptinne et al., 2015). Here, we show that fluctuations also occur at rest in both PAC and waveform shape (Figure 5). Since these measures are correlated, waveform shape metrics may be dynamic during movement as well. This motivates considerations of waveform shape (and PAC) for event-related analyses in future studies. It also indicates that spontaneous fluctuations should be considered for future clinical applications of these signatures as putative physiological biomarkers. Indeed, how each signal is quantified may need to be refined and adapted at the individual patient level. For instance, based on the spread of values over time (Figure 5), we might expect that for some participants very short segments of data would be sufficient to detect a difference in medication state, whereas for others, longer segments would be necessary before an average difference would emerge. For a subset of patients there may not be a clear difference between on and off medication using these measures.

### Uses for a non-invasive electrophysiological biomarker

Non-invasive EEG enables easy and inexpensive measurements in patients and allows for comparisons to healthy controls. Since waveform metrics robustly index medication status using EEG, they can potentially be used as an objective metric reflecting PD state. Waveform shape metrics may be particularly conducive to real-time measures, since they can theoretically be derived from shorter periods of data compared to PAC, for instance. There are many applications for an objective, non-invasive measure of PD state. For instance, such a measure could be used by clinicians to calibrate medications or by patients to determine when to take a medication dose. An objective measure could also be useful for DBS updates, with possible utility for adjusting DBS in real-time (Little et al., 2013; Swann et al., 2018), or for helping find optimal DBS settings, by providing an objective measure which could be used for “automatic programming”. This could optimize the time-consuming and often imprecise process of programming in clinic which usually relies on trial and error assessment. These approaches may be especially useful for PD patients who lack easy access to neurologists, especially those who are mobility impaired or live in remote areas. With increases of wearable technology, one could imagine that acquiring these measures could become increasingly easy even in patients’ own homes

### Limitations

For the on medication recordings, patients had variable amounts of time since their last dose, which could have contributed to some of the variability in our results. However, the fact that differences were still apparent despite this variability, suggests that the findings were robust. Indeed, this variability and associated physiology may be closer to what patients would experience in daily life, reinforcing the translational potential of these approaches. Furthermore, having a small sample size with heterogeneous patients may have added to sample variability. Validation in a larger sample size will be necessary.

Although EEG has an obvious advantage in terms of safety and accessibility compared to invasive recordings, it also has a lower signal-to-noise ratio, poor spatial resolution, and can be impacted by ocular and muscle artifacts. Nevertheless, EEG has greater translational potential and here we demonstrate robust features, even with these methodological shortcomings.

## Conclusion

In PD, basal ganglia thalamocortical loops are excessively synchronized. We show that this synchrony can be detected cortically using scalp EEG as elevated PAC and altered waveform shape in PD patients off medication compared to on. Furthermore, we have shown that considering waveform shape, specifically using a particular shape as an inclusion criterion, increased effect size, suggesting that consideration of waveform shape might be useful for optimizing classification power for non-invasive physiological biomarkers of PD.

## Acknowledgements/Disclosures

We would like to thank Dr. Adam Aron, Jobi George, and the Aron lab for graciously sharing the EEG data set. We would also like to thank Roland Good for statistical consultancy.

S.R.C. is supported by the National Science Foundation Graduate Research Fellowship Program and the University of California, San Diego Chancellor’s Research Excellence Scholarship. B.V. is supported by a Sloan Research Fellowship, the Whitehall Foundation (2017-12-73), and the National Science Foundation (1736028).

N.C.S. is co-inventor on a patent related to the use of PAC recorded with either ECoG or EEG as a signal for closed-loop control of DBS or adjustment of medication dosages.

